# A general model to explain repeated turnovers of sex determination in the Salicaceae

**DOI:** 10.1101/2020.04.11.037556

**Authors:** Wenlu Yang, Zhiyang Zhang, Deyan Wang, Yiling Li, Shaofei Tong, Mengmeng Li, Xu Zhang, Lei Zhang, Liwen Ren, Xinzhi Ma, Ran Zhou, Brian J. Sanderson, Ken Keefover-Ring, Tongming Yin, Lawrence B. Smart, Jianquan Liu, Stephen P. DiFazio, Matthew Olson, Tao Ma

**Affiliations:** Key Laboratory of Bio-Resource and Eco-Environment of Ministry of Education, College of Life Sciences, Sichuan University, Chengdu 610065, China; State Key Laboratory of Grassland Agro-Ecosystem, Institute of Innovation Ecology & College of Life Sciences, Lanzhou University, Lanzhou 730000, China; Department of Biology, West Virginia University, Morgantown, WV 25606, USA; Department of Biological Sciences, Texas Tech University, Lubbock, TX 79409-3131 USA; Departments of Botany and Geography, University of Wisconsin-Madison, 430 Lincoln Dr., Madison, WI 53706, USA; The Key Laboratory of Tree Genetics and Biotechnology of Jiangsu Province and Education Department of China, Nanjing Forestry University, Nanjing, China, 200137; Horticulture Section, School of Integrative Plant Science, Cornell University, New York State Agricultural Experiment Station, Geneva, New York 14456 USA

**Keywords:** Dioecy, Sex determination, Sex chromosome turnover, Genome, *Populus*

## Abstract

Dioecy, the presence of separate sexes on distinct individuals, has evolved repeatedly in multiple plant lineages. However, the specific mechanisms through which sex systems evolve and their commonalities among plant species remain poorly understood. With both XY and ZW sex systems, the family Salicaceae provides a system to uncover the evolutionary forces driving sex chromosome turnovers. In this study, we performed a genome-wide association study to characterize sex determination in two *Populus* species, *P. euphratica* and *P. alba*. Our results reveal an XY system of sex determination on chromosome 14 of *P. euphratica*, and a ZW system on chromosome 19 of *P. alba*. We further assembled the corresponding sex determination regions, and found that their sex chromosome turnovers may be driven by the repeated translocations of a *Helitron*-like transposon. During the translocation, this factor may have captured partial or intact sequences that are orthologous to a type-A cytokinin response regulator gene. Based on results from this and other recently published studies, we hypothesize that this gene may act as a master regulator of sex determination for the entire family. We propose a general model to explain how the XY and ZW sex systems in this family can be determined by the same *RR* gene. Our study provides new insights into the diversification of incipient sex chromosome in flowering plants by showing how transposition and rearrangement of a single gene can control sex in both XY and ZW systems.

## Introduction

The origin and evolution of dioecy (separate sexes) has long been one of the most fascinating topics for biologists (Henry et al., 2018; Feng *et al.*, 2020). The presence of dioecy ensures outcrossing and optimal allocation of reproductive resources for male and female sexual function, thereby providing them with certain advantages in fertility, survival and evolution (Bawa, 1980). In flowering plants, dioecy occurs in only ~6% of all species and has independently evolved thousands of times from hermaphroditic ancestors (Renner and Ricklefs, 1995; Renner, 2014). Many of these species have sex determined by a pair of heteromorphic sex chromosomes that differ in morphology and/or sequence, in the form of male heterogamety (XY system) or female heterogamety (ZW system) (Ming et al., 2011; Charlesworth, 2016). Theory predicts that sex chromosomes evolve from ancestral autosomes via successive mutations in two linked genes with complementary dominance (Charlesworth and Charlesworth, 1978; Charlesworth, 1991). Subsequently, the suppression of recombination between these two sex determination genes progressively spreads along Y or W chromosomes, and permits the accumulation of repetitive elements and duplication or translocation of genomic fragments, which in turn leads to the formation of a sex-specific region and finally degeneration of the sex chromosome (Bergero and Charlesworth, 2009; Charlesworth, 2012; Bachtrog, 2013). Characterizing the genomic architecture of sex in dioecious species is critical for understanding the origin of sex chromosomes, especially in their early stage of evolution.

Over the past decade, impressive progress has been made in unraveling the genetic basis of sex determination in several dioecious plants and the evolutionary history of their sex chromosomes, including papaya (Wang *et al.*, 2012), persimmon (Akagi *et al.*, 2014), asparagus (Harkess *et al.*, 2017), strawberry (Tennessen *et al.*, 2018), date palm (Torres *et al.*, 2018) and kiwifruit (Akagi *et al.*, 2018, 2019). Consistent with the independent origins of sex chromosomes, the sex determination genes identified in these species differ from each another, although most of them function in similar hormone response pathways (Feng *et al.*, 2020). In addition, a recent study found that the sex chromosome turnover in strawberries is driven by repeated translocation of a female-specific sequence (Tennessen *et al.*, 2018). The combined evidence from these studies demonstrates the high variation of plant sex determination mechanisms, and so understanding the factors that drive the convergent evolution of sex chromosomes in plants remains elusive (Zhang *et al.*, 2014).

The family Salicaceae provides an excellent system to study the drivers of sex chromosome evolution. This family includes two sister genera, *Populus* and *Salix*, which are composed exclusively of dioecious species (Peto, 1938; Zhang *et al.*, 2018; Li *et al.*, 2019). Previous studies in multiple *Salix* species have consistently mapped the sex determination regions (SDRs) to chromosome 15, and proposed a ZW system in which females are the heterogametic sex (Pucholt *et al.*, 2015, 2017; Hou *et al.*, 2015; Chen *et al.*, 2016; Zhou *et al.*, 2018, 2020). However, an XY system was recently identified on chromosome 7 in *S. nigra* (Sanderson *et al.*, 2020). In comparison, the SDR has been mapped to multiple locations in different *Populus* species, indicating a dynamic evolutionary history of the sex chromosomes. The SDR has been mapped to the proximal telomeric end of chromosome 19 in *P. trichocarpa* and *P. nigra* (sections *Tacamahaca* and *Aigeiros*) (Gaudet *et al.*, 2007; Yin *et al.*, 2008; Geraldes *et al.*, 2015), and to a pericentromeric region of chromosome 19 in *P. tremula*, *P. tremuloides* and *P. alba* (section *Populus*) (Pakull *et al.*, 2009, 2014; Paolucci *et al.*, 2010; Kersten *et al.*, 2014). Most *Populus* species display an XY sex determination system, but there is some evidence that *P. alba* has a ZW system (Paolucci *et al.*, 2010). Thus far, the only SDR that has been assembled in *Populus* is that of *P. trichocarpa* and *P. deltoides*, and it appears to be much smaller than those observed in *Salix* (Geraldes *et al.*, 2015; Xue *et al.*, 2020). Our recent study on the W chromosome of *S. purpurea* showed intriguing palindromic structures, in which four copies of the gene encoding a type A cytokinin response regulator (*RR*) were identified (Zhou *et al.*, 2020). Interestingly, the ortholog of this gene has also been reported to be associated with sex in *Populus* from section *Tacamahaca* (Geraldes *et al.*, 2015; Bräutigam *et al.*, 2017; Melnikova *et al.*, 2019), which increases the possibility that this gene is an excellent candidate for a common sex determination mechanism in the Salicaceae. However, it is still unclear whether this candidate gene is present in all of these SDRs. Most importantly, how the same gene functions in both the XY and ZW systems remains elusive. Here, we identify the sex determination systems of two additional *Populus* species, *P. euphratica* and *P. alba*, which are from sects. *Turanga* and *Populus* respectively (Wang *et al.*, 2020). We report their complete SDR assemblies and propose a general model to illustrate the potentially shared mechanism of sex determination in this family.

## Results

### Genome assembly

We have previously reported the assembly of the genomes of a male *P. euphratica* (Zhang *et al.*, 2020) and a male *P. alba* var. *pyramidalis* (a variety of *P. alba*) (Ma *et al.*, 2019). Here we further sequenced and *de novo* assembled female genomes for both species using Oxford Nanopore reads. The assembly for the female *P. euphratica* consists of 1,229 contigs with an N50 of 1.7 Mb and a total size of ~529.0 Mb, while the female *P. alba* var. *pyramidalis* assembly has 357 contigs with an N50 of 3.08 Mb, covering a total of ~358.5 Mb (**Table S1**). Both assemblies showed extensive synteny with their respective male reference genomes, and therefore, based on their syntenic relationships, the assembled contigs were anchored onto 19 pseudochromosomes (**Figs. S1 and S2**). The chromosome identities were then assigned by comparison to *P. trichocarpa* (Tuskan *et al.*, 2006).

### XY sex determination on chromosome 14 in *P. euphratica*

In order to characterize the sex determination system of *P. euphratica*, we resequenced the genomes of 30 male and 30 female individuals (**Table S2**) and performed a genome-wide association study (GWAS). Using the male assembly as the reference genome, a total of 24,651,023 high-quality single nucleotide polymorphisms (SNPs) were identified. After Bonferroni correction, we recovered 310 SNPs significantly associated with sex (α<0.05; **Figs. 1A**, **S3A and Table S3**). In-depth analysis found that almost all genotypes (99.99%) of these sex-associated loci are homozygous in females, while 93.57% of the genotypes are heterozygous in males (**Fig. 1B**). A similar pattern was observed when the sex association analysis was performed by using the female assembly as the reference genome (**Figs. S3B and S4, and Tables S4 and S5**). These results consistently indicate that an XY system is involved in sex determination of *P. euphratica*.

**Fig. 1.**
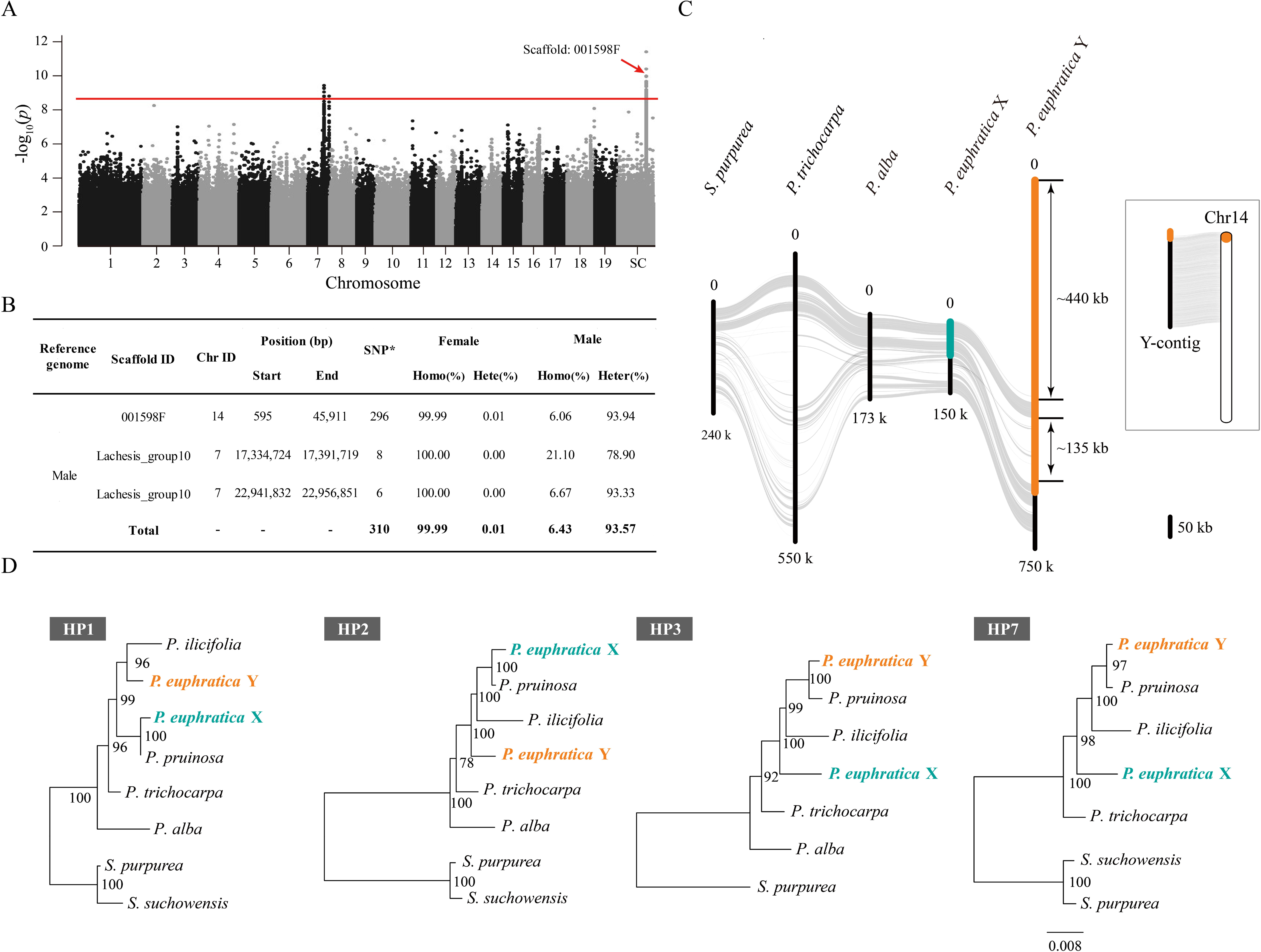
XY sex determination on chromosome 14 in *P. euphratica*. **(A)** Manhattan plot of *P. euphratica* based on the results of genome-wide association study (GWAS) with the male genome as reference. The y-axis represents the strength of association (−log_10_(*P* value)) for each SNP sorted by chromosomes and scaffolds (SC; x-axis). The red line indicates the significance after Bonferroni multiple corrections (α < 0.05). Note that the scaffold ‘001598F’ is located on chromosome 14 based on its syntenic relationship with the proximal end of chromosome 14 of *P. trichocarpa*. (**B**) Summary of male *P. euphratica* genome regions containing SNPs significantly associated with sex. SNP*, significantly associated SNPs; Homo, Homozygous; Hete, Heterozygosis. **(C)** Synteny relationships between our assembled Y-contig and X chromosome of *P. euphratica*, as well as the corresponding region of chromosome 14 for *P. alba*, *P. trichocarpa* and *S. purpurea*. The highlighted part represents the sex determination region (SDR), yellow for Y-SDR and green for X-SDR. Schematic diagram showing the corresponding position of the SDR on chromosome 14 of *P. euphratica*. **(D)** Phylogenetic relationships of the homolog pairs (HP) shared between Y- and X-SDR of *P. euphratica* and their orthologous genes in other Salicaceae species. Detailed information about these genes is listed in Table S7 and additional phylogenetic trees are shown in Fig. S7. Note that only the orthologous genes located on the corresponding region of chromosome 14 were used for phylogenetic analysis.

In addition, we found that the vast majority of the significantly sex-associated SNPs were located at the proximal end of chromosome 14 (the un-anchored scaffold ‘001598F’ in male genome was located onto chromosome 14 based on its syntenic relationship with *P. trichocarpa* genome), while a few other SNPs were present at chromosomes 7, 9, 12 and 19 (**Figs. 1A, 1B and S4, and Table S5**). We then attempted to use ultra-long nanopore reads generated from a male individual (**Table S6**) to further reconstruct a new assembly with X and Y haplotypes as separate contigs. This led to the identification of a contig that was highly similar to the sex-associated regions and specifically contained Y-linked alleles (**Fig. S5**). The Y-linked region was further determined by examining the relative depth of coverage when aligning male versus female resequencing reads against the reference (**Fig. S6**). Based on the syntenic relationship, the SDR of *P. euphratica* can be mapped to the proximal end of chromosome 14 and the Y-linked region is about 658 kb in length, corresponding to ~84 kb on the X chromosome (**Fig. 1C**). We found that two segments spanning 440 kb and 135 kb respectively, are specific to the Y-linked region (**Fig. 1C**), suggesting the occurrence of significant chromosome divergence between the X and Y haplotypes, which can be maintained by suppressed recombination.

We predicted a total of 37 protein-coding genes in the Y-linked region, many of which have high similarity with genes on other autosomes and are considered as translocated genes (**Table S7**). Among these, we found that 9 of the Y-specific genes were annotated as members of the LONELY GUY (LOG) family, which encodes cytokinin-activating enzymes that play a dominant role in the maintenance of the shoot apical meristem and in the establishment of determinate floral meristems (Kuroha *et al.*, 2009; Tokunaga *et al.*, 2012; Han and Jiao, 2015). Ten genes were identified in both X and Y haplotypes. A phylogenetic analysis of these genes showed that the X and Y alleles began to diverge after their split with *P. trichocarpa* and *P. alba* (**Figs. 1D and S7**), suggesting that the SDR of *P. euphratica* appears to be established relatively recently.

### ZW sex determination on chromosome 19 in *P. alba*

We used a similar GWAS strategy for 30 male and 30 female resequenced individuals to characterize the sex determination system of *P. alba* (**Table S8**). When the male and female assembly was used as a reference genome, respectively, 173 and 55 SNPs that were significantly associated with sex were identified (**Figs. 2A, 2B, S8 and S9, and Tables S9-S11**). Most of the sex-associated SNPs are heterozygous in females and homozygous in males (**Fig. 2B and Table S10**), confirming the ZW sex determination system in *P. alba*, which was also suggested based on genetic mapping in a previous study (Paolucci *et al.*, 2010).

**Fig. 2.**
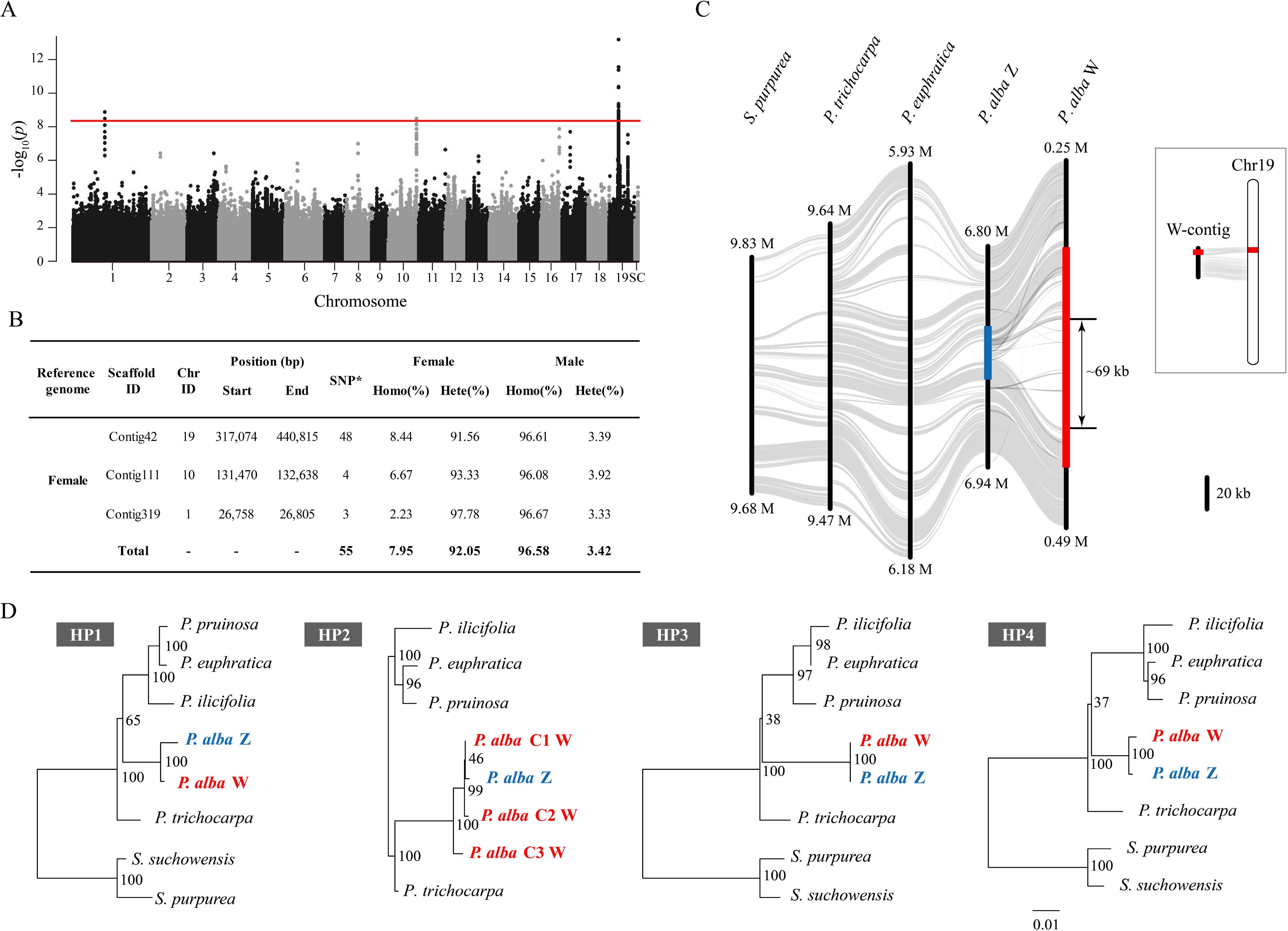
ZW sex determination on chromosome 19 in *P. alba*. **(A)** Manhattan plot of *P. alba* based on the results of GWAS with female genome as reference respectively. The y-axis represents the strength of association (−log_10_(*P* value)) for each SNP sorted by chromosomes and scaffolds (SC; x-axis). The red line indicates the significance after Bonferroni multiple corrections (α < 0.05). (**B**) Summary of female *P. alba* genome regions containing SNPs significantly associated with sex. SNP*, significantly associated SNPs; Homo, Homozygous; Hete, Heterozygosis. **(C)** Synteny relationships between our assembled W-contig and Z chromosome of *P. alba*, as well as the corresponding region of chromosome 19 for *P. euphratica*, *P. trichocarpa* and *S. purpurea*. The highlighted part represents SDR, red for W-SDR and blue for Z-SDR. Schematic diagram showing the corresponding position of the SDR on chromosome 19 of *P. alba*. **(D)** Phylogenetic relationships of the homolog pairs (HP) shared between W- and Z-SDR of *P. alba* and their orthologous genes in other Salicaceae species. The detail information of these genes is listed in Table S12. Note that there are 3 copies for ‘HP2’ on the W-SDR of *P. alba*, and only the orthologous genes located on the corresponding region of chromosome 19 were used for phylogenetic analysis.

We found that these sex-associated SNPs are mainly located on a non-terminal region of chromosome 19 (**Figs. 2A, 2B and S8, and Table S10**). Next, we examined the female-specific depth profile, combined with the support of ultra-long nanopore reads (**Table S6**), to delineate the W haplotype of *P. alba* to a region of about 140 kb on chromosome 19, with a corresponding Z haplotype that is only 33 kb in length (**Figs. 2C, S10 and S11**). Compared to the Z haplotype and corresponding autosomal regions of the other Salicaceae species, a specific insertion of 69 kb was observed in the W haplotype, indicating a recent origin of the SDR in *P. alba*.

Sequence annotation predicted 18 protein-coding genes in the W haplotype, six of which were also found in the Z haplotype (**Table S12**). The high identity of these alleles between the W and Z haplotype suggests that recombination suppression occurred very recently (**Fig. 2D**). We further found that the gene encoding NAC-domain protein, *SOMBRERO* (*SMB*), which has a similar function to the *VND*/*NST* transcription factors that regulate secondary cell wall thickening in woody tissues and maturing anthers of *Arabidopsis* (Mitsuda *et al.*, 2005; Bennett *et al.*, 2010), was expanded from one member in the Z haplotype to three copies in the W haplotype (‘HP2’ in **Fig. 2D**). There are 12 genes specific to the W haplotype (**Table S12**), including *DM2H* (*DANGEROUS MIX2H*), which encodes a nucleotide-binding domain and leucine-rich repeat immune receptor protein (Chae *et al.*, 2014); *CCR2* (Cinnamoyl CoA reductase), which is involved in lignin biosynthesis and plant development (Thevenin *et al.*, 2011); and *STRS1* (*STRESS RESPONSE SUPPRESSOR1*), a gene encoding a DEAD-box RNA helicase, which is involved in epigenetic gene silencing related to stress responses (Khan *et al.*, 2014). More interesting, we also identified three copies of the gene encoding a type A cytokinin response regulator (*RR*) in the W-specific region (**Fig. 3A**), the ortholog of which has also been identified to be associated with sex determination in poplar and willow (Geraldes *et al.*, 2015; Bräutigam *et al.*, 2017; Melnikova *et al.*, 2019; Zhou *et al.*, 2020). Very little sequence differences were found among these three copies, and combined with the fact that the ortholog of the *RR* gene is located at the distal end of chromosome 19 in *P. trichocarpa* and *P. euphratica* (**Fig. 3**), we conclude that the *RR* gene was translocated from the end of chromosome 19 to the W haplotype of *P. alba* and then underwent at least two rounds of recent duplication.

**Fig. 3.**
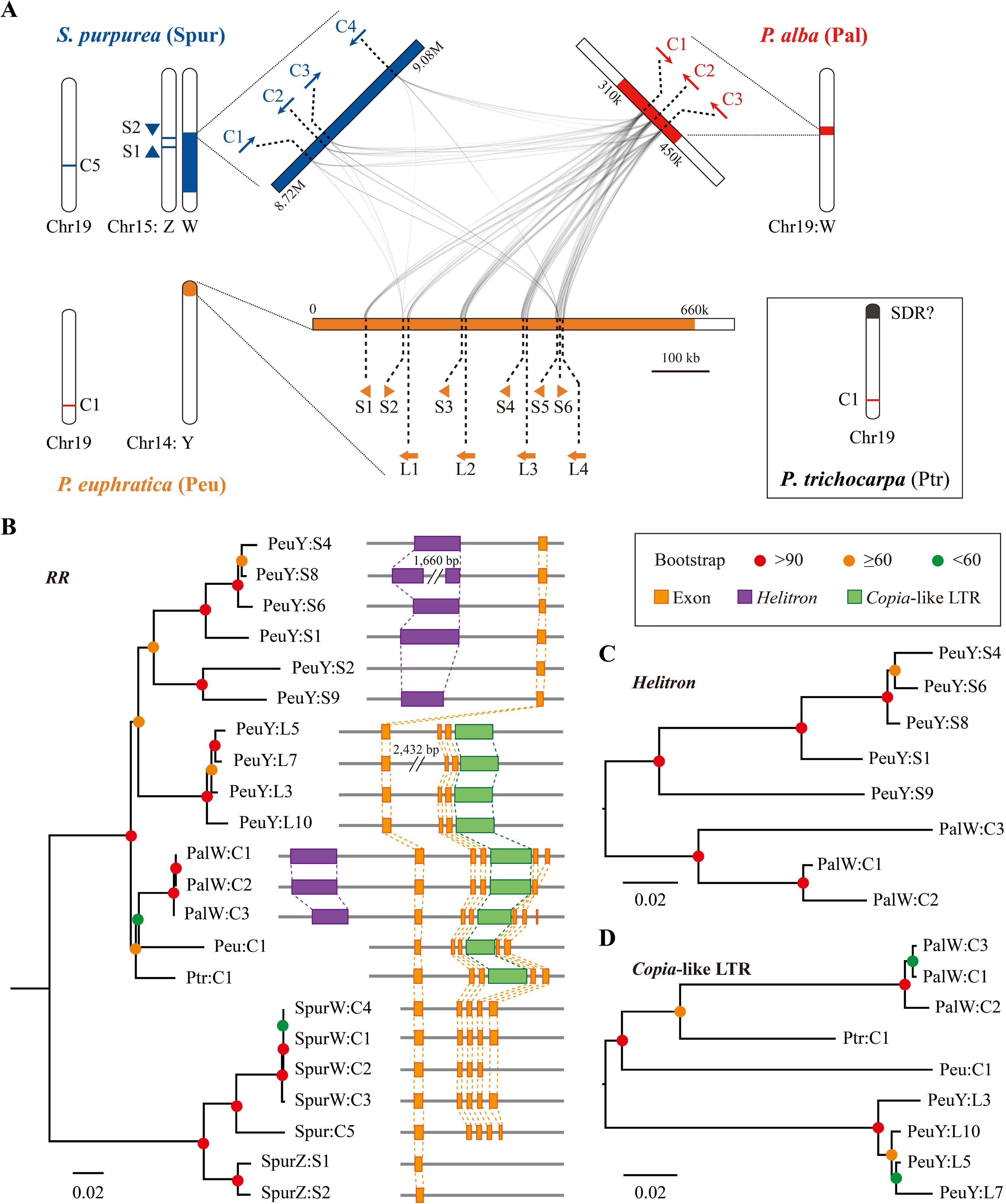
Evidence for SDR turnover in Salicaceae. **(A)** Synteny relationships among the Y-SDR of *P. euphratica* (yellow) and the W-SDRs of *P. alba* (red) and *S. purpurea* (blue), showing the copies of *RR* intact gene (‘C’) and partial duplicates (‘S’: small duplicate; ‘L’: large duplicate) on their SDRs. For each species, corresponding positions for other *RR* gene copies or partial duplicates on the autosome are also shown. **(B)** Phylogenetic relationship of the *RR* sequences (including intact genes and partial duplicates) identified in the four species. The tree was rooted by a paralogous gene ‘*RR16*’. The gene structures and relative positions of *Helitron* and *Copia*-like LTR are also shown. Phylogenetic relationships of the *Helitron* **(C)** and *Copia*-like LTR **(D)** around the *RR* sequences. All the sequences were named according to Fig. 3A. Peu: *P. euphratica*; Pal: *P. alba*; Ptr: *P. trichocarpa*; Spur: *S. purpurea*.

### Evidence for SDR turnover in Salicaceae

We have shown that *P. euphratica* and *P. alba* have different sex determination systems, and that the SDRs are different from those reported in *P. trichocarpa* and *S. purpurea*, indicating extraordinarily high diversity of sex determination in the Salicaceae. In order to examine whether the sex determination regions originated independently in each lineage, or evolved into the current SDRs separately after a common ancient origin, we performed syntenic analysis on these SDRs in *P. euphratica* and *P. alba*, and the corresponding autosomal regions in *P. trichocarpa* and *S. purpurea*. We found that although the pseudo-autosomal regions of these sex chromosomes are highly collinear with their corresponding autosomal regions in other species, the sequences in the sex-specific regions are not alignable (**Figs. 1C and 2C**). In contrast, although there was little collinearity among these SDRs, a homologous sequence with multiple duplicates was identified between the Y haplotype of *P. euphratica* and the W haplotype of *P. alba* **(Fig. 3A)**. Interestingly, the locations of the duplicates overlapped with the three predicted *RR* genes in *P. alba*. In the corresponding regions of the Y haplotype of *P. euphratica*, we identified 10 partial duplicates of the RR gene including four covering the first three exons (large duplicate) and six covering only the first exon (small duplicate) of the *RR* gene (**Fig. 3**). Phylogenetic analysis of these duplicates showed that the three *RR* genes in *P. alba* clustered together and are closely related to the intact orthologs of *P. euphratica* and *P. trichocarpa*, while the partial duplicates from *P. euphratica* divided into two main clades, one with only large duplicates and a second clade with only small duplicates (**Fig. 3B**).

Since the *RR* duplicates were found in the SDRs of all of the current and previously studied species, we believe that they may play important roles in sex determination of the Salicaceae species. These results also lead to the hypothesis that these species shared an ancient origin of sex chromosomes, followed by frequent turnover events due to translocation of the *RR* duplicates. This is further supported by the distant relationship between the partial and intact *RR* duplicates **(Fig. 3B)**, which indicate that the partial duplicates originated before the divergence of these poplar species and were repeatedly inserted into the SDRs of *P. euphratica*. We did not detect any structurally intact long terminal repeat retrotransposons (LTR-RTs) around these *RR* duplicates, which made it impossible to estimate their insertion time. However, around the *RR* duplicates in *P. euphratica*, we identified a *Helitron*-like transposable element upstream of each small duplicate except the second one (‘PeuY:S2’), and a *Copia*-like LTR fragment in the downstream region of each large duplicate **(Fig. 3B)**. These two repetitive elements were also identified in all three *RR* duplicates of *P. alba*, and are located upstream and in the third intron of the *RR* gene, respectively, similar to that in *P. euphratica*. The phylogenetic trees of the two elements and the *RR* duplicates exhibited a similar topological relationship, suggesting that they may be transposed together as a unit (**Figs. 3C and 3D**). The extremely high similarity of these sequences indicates that they were recently transposed into the SDRs of *P. euphratica* and *P. alba*, respectively, consistent with the observation that their sex chromosomes have not been severely degenerated. In addition, we found that the *Helitron*-like element was not present in the upstream region of the intact *RR* genes at chromosome 19 of *P. euphratica* and *P. trichocarpa* (**Fig. 3B**), which led us to speculate that this element may be the main driving force for gene replication during the evolution of SDRs in *P. euphratica* and *P. alba*. However, we failed to detect the same pattern in *S. purpurea*, in which multiple *Copia* LTR-RTs were predicted instead of the *Helitron* elements (Zhou *et al.*, 2020). This implies that poplar and willow may have different SDR turnover mechanisms, which requires further evidence from more species to confirm.

## Discussion

It is notoriously difficult to assemble the complete sequence of SDRs or sex chromosomes, which usually have a high repeat density and many translocated segments from autosomes (Charlesworth, 2012; Bachtrog, 2013). In our study, the sex-associated loci were initially mapped onto multiple different chromosomes **(Figs. 1 and 2)**, although they consistently revealed an XY sex determination system in *P. euphratica* and a ZW system in *P. alba*. These results may be caused by the lack and/or mis-assembly of SDRs in the reference genome, especially when the genome from a homozygous (XX or ZZ) individual was used as reference, the reads from Y- or W-specific regions of hemizygous (XY or ZW) individuals may be misaligned to homologous sequences on autosomes and led to false associations. Similar phenomena were also observed in the sex association analysis of *P. trichocarpa*, *P. balsamifera* and *S. purpurea*, which may lead to an inaccurate localization of SDRs in assemblies (Geraldes *et al.*, 2015; Zhou *et al.*, 2020). The high sequence similarity between these sex-associated regions and the SDRs we finally established strongly supports this possibility **(Figs. S5 and S10**). Therefore, our research emphasizes the importance and necessity for precise assembly of SDRs using multiple complementary methods, including the ultra-long read sequencing, haplotype phased assembly and the sex-specific depth of read mapping.

Our results further indicate that the SDRs of poplar species are generally shorter in length and contain relatively fewer genes than that recently reported in *S. purpurea* (Zhou *et al.*, 2020), though the size of this SDR may be inflated due to overlap with the centromere (Zhou *et al.*, 2018). Although some specific insertions were observed on the Y and W chromosomes, we found no obvious degeneration of sex chromosomes at least in *P. euphratica* and *P. alba*. These results suggest that the SDRs of these two species were established relatively recently, which is a common feature of the sex chromosomes of the Salicaceae species studied so far (Geraldes *et al.*, 2015; Pucholt *et al.*, 2017; Zhou *et al.*, 2018, 2020). Along with this, our results also suggest that the Y and W chromosomes have expanded in content, a pattern that is common in young sex chromosomes of plants (Hobza *et al.*, 2015, 2017). Moreover, our results simultaneously showed that the Salicaceae exhibit an extremely fast rate of sex-chromosome turnover. In previous studies, SDRs have been reported only on chromosome 15 with female heterogamety (ZW) in willow except *S. nigra* (Pucholt *et al.*, 2015, 2017; Hou *et al.*, 2015; Chen *et al.*, 2016; Zhou *et al.*, 2018, 2020; Sanderson *et al.*, 2020), and on chromosome 19 of poplar with most species showing male heterogamety (XY) (Gaudet *et al.*, 2007; Yin *et al.*, 2008; Pakull *et al.*, 2014; Geraldes *et al.*, 2015). However, our study identified an XY system with the SDR on chromosome 14 of *P. euphratica* for the first time, and confirmed a ZW system with SDR on chromosome 19 of *P. alba*. These results highlight the complexity and diversity of sex determination in this family. Comparative analysis showed that translocation of genes from autosomes to the SDR and gene replication frequently occurred both on the Y chromosomes of *P. euphratica* and on the W chromosomes of *P. alba*, indicating that these two events are likely to be important contributors during SDR turnover. The regulatory mechanisms and functions of these genes in sex determination and sexual dimorphism in these two species need further investigation.

Among all genes on SDRs, the cytokinin response regulator is the most likely candidate for controlling sex determination in the Salicaceae, not only because the orthologs of this gene have been found to be sex-associated in most of the reported species in the family, but also because it is the only homologous sequence found in the sex chromosomes of *P. euphratica*, *P. alba*, *P. trichocarpa*, *P. deltoides* and *S. purpurea* (**Fig. 3**), the only Salicaceae species with SDR precisely assembled (Zhou *et al.*, 2020; Xue *et al.*, 2020). Recent progress has revealed that the genes involved in cytokinin signaling play important roles in the regulation of unisexual flower development in plants (Wybouw *et al.*, 2019; Kieber *et al.*, 2018; Feng *et al.*, 2020). Specifically, a Y-specific type-C cytokinin response regulator (*Shy Girl*, *SyGI*) was recently identified as a suppressor of carpel development and therefore is a strong candidate of sex determination in kiwifruit (Akagi *et al.*, 2018). Similar to the pattern of the *RR* genes found in the Salicaceae species, in kiwifruit *SyGI* was duplicated from an autosome and subsequently gained a new function on its Y chromosome. However, the type-A *RR* genes we identified here are not orthologous to the *SyGI* gene, so we speculate that they may have different functions in the cytokinin signaling pathway. Based on our results, it is reasonable to suspect that the *RR* genes are more likely to function as a dominate promoter of female function **(Fig. 4)**, as they exist on the W chromosomes of both *P. alba* and *S. purpurea* in intact duplicates. In contrast, the RR gene fragments on the Y chromosome of *P. euphratica* exist as two partial duplicates with different sizes. This may serve as a female suppressor by encoding an siRNA that targets the intact *RR* gene at the distal end of chromosome 19, possibly through RNA-directed DNA methylation (Brautigam *et al.*, 2017; Xue *et al.*, 2020). It should be noted that, although the intact *RR* gene has been reported to be associated with sex in *P. trichocarpa*, there is still no evidence to support the gene’s localization on its Y chromosome. In the previous GWAS study (Geraldes *et al.*, 2015), most of the sex-associated loci of *P. trichocarpa* were located on the proximal end of chromosome 19. The associated signals scattered around the intact *RR* gene, which is located at the distal end of chromosome 19, were most likely due to assembly errors arising from the fact that this reference genome is derived from a female (XX) individual (the major factor in misleading SDR localization as mentioned above). Therefore, our findings consistently showed that Salicaceae species potentially share a common mechanism of sex determination, in which the specific duplication of the *RR* orthologs on SDRs may have played an important role in the acquisition of separate sexes in these species.

**Fig. 4.**
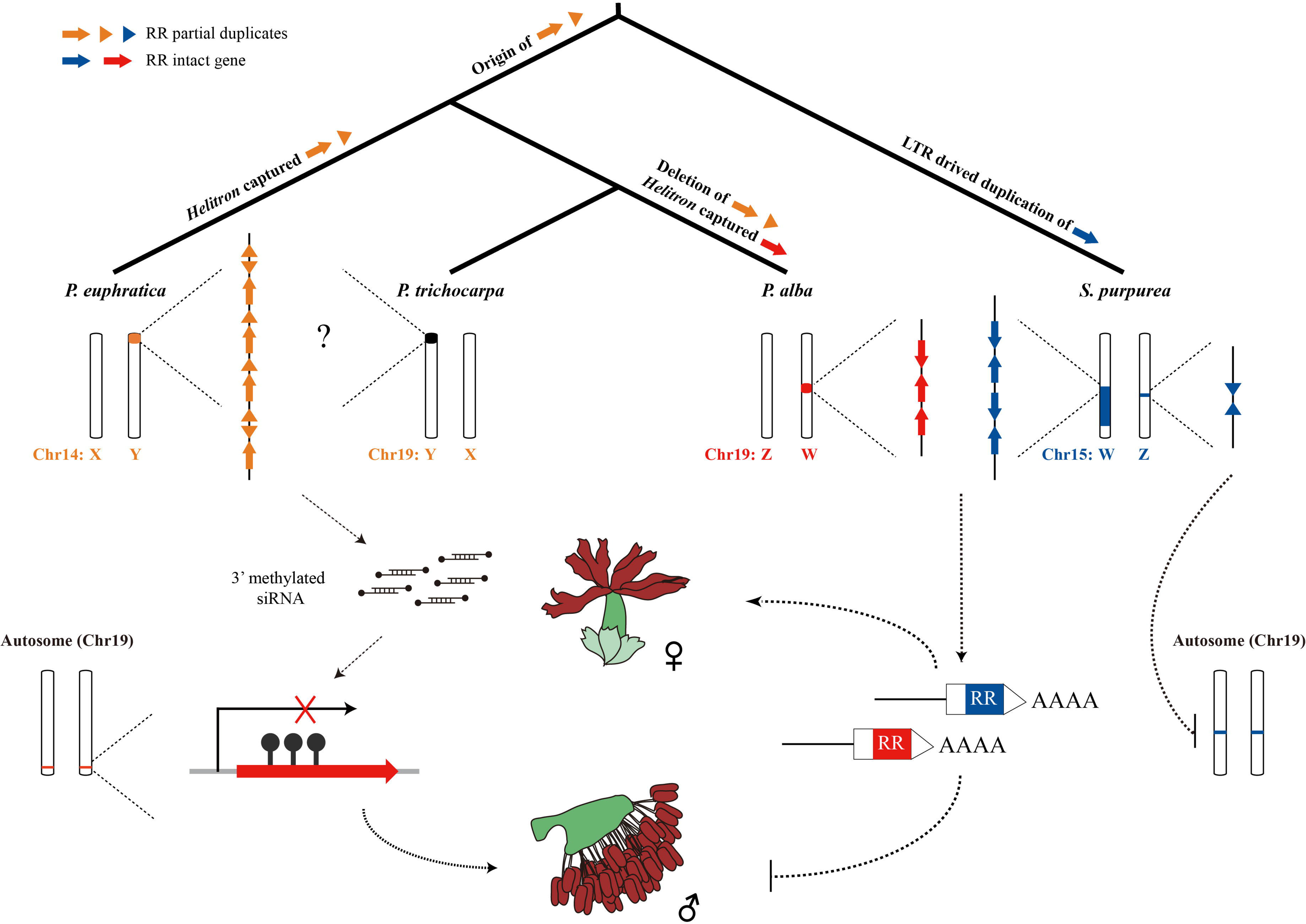
Hypothetical model for sex system turnovers in Salicaceae. The W chromosomes of *P. alba* and *S. purpurea* both carry several intact *RR* genes and are likely to serve as a dominate promoter of female function. On the Y chromosome of *P. euphratica*, partial duplicates of the *RR* gene are like to serve as a female suppressor by encoding an siRNA that targets the intact RR gene through RNA-directed DNA methylation. Note that Y-SDR of *P. trichocarpa* has not yet been assembled, so whether a similar pattern should be found in this species remains to be confirmed.

More interestingly, we identified *Helitron*-like repetitive elements upstream of the *RR* duplicate in both SDRs of *P. euphratica* and *P. alba*, regardless of whether the *RR* duplicate is intact or partial (**Fig. 3**). As a major class of DNA transposons, *Helitrons* were hypothesized to transpose by a rolling circle replication mechanism, and have been found to frequently capture genes or gene fragments and move them around the genome, which is believed to be important in the evolution of host genomes (Morgante *et al.*, 2005; Kapitonov and Jurka, 2007). Our results suggest that the *RR* fragments and intact gene sequences appear to have been captured by *Helitrons* in *P. euphratica* and *P. alba*, and subsequently replicated in their SDRs **(Figs. 3 and 4)**. Furthermore, our phylogenetic analysis indicated that the intact *RR* gene was captured very recently in *P. alba*, at least after its split with *P. trichocarpa* (**Fig. 3B**). In contrast, although we found high similarity among the *RR* partial duplicates of *P. euphratica*, these sequences are quite different from the intact *RR* genes of other poplar species **(Fig. 3B)**. These results indicate that the partial duplicates were present before the diversification of poplar species, but only recently replicated on the Y chromosome of *P. euphratica*. We found that the partial duplicate of the *RR* gene is lacking in *P. alba*, which may be another key event in addition to the duplication of the intact *RR* gene, in the transition of the sex determination system from XY to ZW **(Fig. 4)**. In addition, the high nucleotide identity among intact *RR* genes of *S. purpurea* reflects another possible SDR turnover event in willow, which might be driven by the replication of a *Copia* LTR (Zhou *et al.*, 2020), rather than by a *Helitron* as we found in poplar. Moreover, we also identified an inverted repeat of the first exon of the *RR* gene and an intact copy on the chromosomes 15Z and 19 of *S. purpurea*, respectively **(Fig. 3)**. This suggests a model whereby the inverted repeat is suppressing the *RR* gene of chromosome 19 in males, but the SDR on the W chromosome may be dominant to this effect in females, possibly due to increased dosage or another mechanism **(Fig. 4)**. These observations further indicate that the sex determination system of *S. purpurea* may have been changed from XY to ZW relatively recently, since the suppressing mechanism from the *RR* partial duplication is still retained. This turnover was also supported by the XY sex determination system of the basal *Salix* species, *S. nigra* (Sanderson *et al.*, 2020). Therefore, our results suggest that the high activity of these repetitive elements is the most likely cause of the recently established SDRs in these species, and further indicate that at least three turnover events have occurred in the evolution of sex chromosomes of the Salicaceae species **(Fig. 4).**

In conclusion, here we present an XY system of sex determination with the SDR on the proximal end of chromosome 14 in *P. euphratica*, and a ZW system with the SDR on a non-terminal region of chromosome 19 in *P. alba*. Both SDRs appear to have evolved relatively recently and are characterized by frequent translocations from autosomes and gene replication events. Our comparative analysis also demonstrated an extremely fast rate of sex chromosome turnover among Salicaceae species, which may be driven by *Helitron* transposons in poplar and by *Copia* LTRs in willow. Most importantly, we propose a model showing that poplar and willow have a common underlying mechanism of sex determination, which controls the XY and ZW systems simultaneously through a type-A *RR* gene. In the future, it will be necessary to conduct transgenic function experiments and comparative analysis from more species in this family to further support our model.

## Methods

### Genome sequencing

We have previously reported the reference genome of a male *P. euphratica* (Zhang *et al.*, 2020) and a male *P. alba* (Ma *et al.*, 2019). In this study, we further collected the fresh leaves of a female *P. euphratica* and a female *P. alba* for genome sequencing and assembly. Genomic DNA was extracted using the QIAGEN Genomic DNA extraction kit (Qiagen, Hilden, Germany) following the manufacturer’s protocol. To generate Oxford Nanopore long reads, approximately 15 μg of genomic DNA was size-selected using the BluePippin system (Sage Science, USA), and processed according to the protocol of Ligation Sequencing Kit (SQK-LSK109). The final library was sequenced on a PromethION sequencer (Oxford Nanopore Technologies, UK) with a running time of 48 hours. The Oxford Nanopore proprietary base-caller, Albacore v2.1.3, was used to perform base calling of the raw signal data and convert the FAST5 files into FASTQ files.

In addition, paired-end libraries with insert size of ~300 bp were also constructed using NEB Next® Ultra DNA Library Prep Kit (NEB, USA), with the standard protocol provided by Illumina (San Diego, CA, USA). The library was sequenced on an Illumina HiSeq X Ten platform (Illumina, San Diego, CA, USA). These sequencing data were used for correction of errors inherent to long read data for genome assembly.

### Genome Assembly

For genome assembly, we first removed the Nanopore long reads shorter than 1 kb and the low-quality reads with a mean quality ≤ 7. The long reads underwent self-correction using the module ‘NextCorrect’ and then assembled into contigs using ‘NextGraph’ implemented in Nextdenovo v2.2.0 (https://github.com/Nextomics/NextDenovo) with default parameters. Subsequently, the filtered Nanopore reads were mapped to the initial assembly using the program Minimap2 v2.17-r941 (Li, 2018) and NextPolish v1.0 (https://github.com/Nextomics/NextPolish) was used with three iterations to polish the genome. In addition, we further aligned the Illumina reads to the genome using BWA-MEM v0.7.15 (Li and Durbin 2009) and corrected base-calling by an additional three rounds of NextPolish runs with default parameters. Finally, the corrected genome was aligned to their respective male reference genome using the LAST program (Kielbasa *et al.*, 2011) and the syntenic relationships were used to anchor the assembled contigs onto 19 chromosomes.

### Population sample collection, resequencing and mapping

Silica gel dried leaves of *P. euphratica* and *P. alba* were collected from wild populations in western China. For each species, the sex of 30 male and 30 female individuals was identified from flowering catkins. Genomic DNA of each sample was extracted using the Qiagen DNeasy Plant Minikit (Qiagen, Hilden, Germany). Paired-end libraries were prepared using the NEBNext Ultra DNA Library Prep Kit (NEB, USA) and sequenced on an Illumina HiSeq X Ten platform, according to the manufacturer’s instructions.

The generated raw reads were first subjected to quality control and low-quality reads were removed if they met either of the following criteria (Ma *et al.*, 2018): i) ≥10% unidentified nucleotides (N); ii) a phred quality ≤ 7 for > 65% of read length; iii) reads overlapping more than 10 bp with the adapter sequence, allowing < 2 bp mismatch. Reads shorter than 45 bp after trimming were also discarded. The obtained high-quality cleaned reads were subsequently mapped to the male and female reference genomes of each species, respectively, using BWA-MEM v0.7.15 with default parameters (Li and Durbin 2009). Then the alignment results and marked duplicate reads were sorted using SAMtools v0.1.19 (Li *et al.*, 2009). Finally, Genome Analysis Toolkit (GATK) (DePristo *et al.*, 2011) was performed to process base quality recalibrations to enhance alignments in regions around putative indels with two steps: i) ‘RealignerTargetCreator’ was applied to identify regions where realignment was needed; ii) ‘IndelRealigner’ was used to realign these regions.

### SNP calling, filtering and genome-wide association study (GWAS)

To prevent biases in SNP calling accuracy due to the difference of samples size between groups, single-sample SNP and genotype calling were first implemented using GATK (DePristo *et al.*, 2011) with ‘HaplotypeCaller’, and then multi-sample SNPs were identified after merging the results of each individual by ‘GenotypeGVCFs’. A series of filtering steps were performed to reduce false positives (Yang *et al.*, 2018), including removal of (1) indels with a quality scores < 30, (2) SNPs with more than two alleles, (3) SNPs at or within 5 bp from any indels, (4) SNPs with a genotyping quality scores (GQ) < 10, and (5) SNPs with extremely low (< one-third average depth) or extremely high (> threefold average depth) coverage. The identified SNPs were used for subsequent GWAS analysis. A standard case/control model between allele frequencies and sex phenotype was performed using Plink v1.9 (Purcell *et al.*, 2007). For each species, associations at α < 0.05 after Bonferroni correction for multiple testing were reported as the significantly sex-associated SNPs. These sex-associated SNPs that occurred within 10 kb on the same chromosome were merged into the same interval.

### Construction of *P. euphratica* Y contig and *P. alba* W contig

To construct the Y contig of *P. euphratica* and the W contig of *P. alba*, we further generated ultra-long sequences from a male (XY) *P. euphratica* and a female (ZW) *P. alba*, using an optimized DNA extraction followed by modified library preparation based on the Nanopore PromethION sequencer (Jain *et al.*, 2018; Gong *et al.*, 2019). For *P. euphratica*, we did not find contigs that clearly contained Y-linked sequences in its male genome, which may be due to assembly errors, so we used multiple methods to determine its Y contig. At first, we attempted to find the male-specific k-mers from the high-quality resequencing reads of both male and female samples. Briefly, all 32 bp k-mers starting with the ‘AG’ dinucleotide were extracted from all resequencing reads, and the number of occurrences of each specific subsequence in female and male individuals was counted, respectively. The use of the ‘AG’ dinucleotide is to reduce the number of k-mer sequences and effectively speed up the analysis. The k-mer counts were then compared between male and female, and the male-specific k-mers (female count was 0) were obtained. Next, we extracted the ultra-long nanopore reads containing at least one of the identified male-specific k-mers, and assembled these ultra-long reads using the software Canu v1.7 (Koren *et al.*, 2017), resulting in a ‘male-specific contig’ that was 450 kb in length. Simultaneously, we also *de novo* assembled all of the ultra-long nanopore reads into a draft male genome using Nextdenovo v2.2.0. By comparing the ‘male-specific contig’ with the obtained male genome, we identified a candidate Y contig that contained a large number of male-specific alleles and exhibited a widespread synteny and continuity with the ‘male-specific contig’. To further refine the sex determination region along this candidate Y contig, we re-mapped the resequencing data to the draft genome by BWA-MEM v0.7.15 (Li and Durbin, 2009), and extracted the average depth of coverage using a non-overlapping sliding window (1 kb in length) by SAMtools v0.1.19 (Li *et al.*, 2009). Finally, we compared the relative depth of coverage between male and female individuals, and found that the region between 0 and 658 kb of this contig showed male-specific depth and was therefore considered to be the sex determination region on the Y chromosome of *P. euphratica*.

For *P. alba*, we first performed a whole genome alignment between its male and female genome using the program LAST (Kielbasa *et al.*, 2011). Fortunately, we found that the sex-associated region in the female genome contained a large insert compared to the corresponding region in the male genome. We used the same method as above to count the relative depth of coverage between male and female individuals of *P. alba*, and found that the region between 310 and 450 kb of this contig exhibited female-specific depth. Therefore, this region was directly considered to be the sex determination region on the W chromosome of *P. alba*, and the assembly accuracy of this region was also confirmed by our ultra-long nanopore reads.

### Annotation and comparison of the Y and W contigs

Transposable elements in our assembled Y and W contigs were identified and classified using the software RepeatMasker (Tarailo-Graovac and Chen, 2009). Gene annotation was conducted by combining the results of *de novo* prediction from the program Augustus v.3.2.1 (Stanke *et al.*, 2006), homology-based prediction using the protein sequences of *A. thaliana*, *P. trichocarpa* and *S. purpurea* downloaded from Phytozome 12 (https://phytozome.jgi.doe.gov/), as well as transcriptome data of *P. euphratica* and *P. alba* generated from our previously studies (Ma *et al.*, 2019; Hu *et al.*, 2020; Zhang *et al.*, 2020). The predicted genes were searched against predicted proteins from *P. trichocarpa*, *S. suchowensis* and *A. thaliana* to find the closest homologous annotation.

To construct the phylogenetic relationships among the allelic genes on the X/Y or Z/W contigs, we further identified their orthologous genes in *P. pruinosa* (Yang *et al.*, 2017), *P. ilicifolia* (Chen *et al.*, 2020) and *S. suchowensis* (Dai *et al.*, 2014) genomes by combining reciprocal blast results and their syntenic relationships. The sequences were aligned using ClustalW with default parameters provided in MEGA5 (Tamura *et al.*, 2011) and the resulting alignments were adjusted manually. A maximum likelihood tree was built using MEGA5 with default parameters.

## Accession numbers

The whole genome sequence data reported in this paper have been deposited in the Genome Warehouse in BIG Data Center (BIG Data Center Members, 2019), Beijing Institute of Genomics (BIG), Chinese Academy of Sciences, under accession number PRJCA002485 that is publicly accessible at https://bigd.big.ac.cn/bioproject.

## Acknowledgements

This research was supported by National Natural Science Foundation of China (31561123001, 31922061, 41871044, 31500502), NSF Dimensions of Biodiversity Program (1542509 to S.D. and 1542599 to M.O.), National Key Research and Development Program of China (2016YFD0600101), Fundamental Research Funds for the Central Universities (SCU2019D013).

## References

Akagi, T., Henry, I.M., Ohtani, H., Morimoto, T., Beppu, K., Kataoka, I., and Tao, R. (2018). A Y-Encoded Suppressor of Feminization Arose via Lineage-Specific Duplication of a Cytokinin Response Regulator in Kiwifruit. The Plant cell 30:780–795.

Akagi, T., Henry, I.M., Tao, R., and Comai, L. (2014). Plant genetics. A Y-chromosome-encoded small RNA acts as a sex determinant in persimmons. Science 346:646–650.

Akagi, T., Pilkington, S.M., Varkonyi-Gasic, E., Henry, I.M., Sugano, S.S., Sonoda, M., Firl, A., McNeilage, M.A., Douglas, M.J., Wang, T., et al. (2019). Two Y-chromosome-encoded genes determine sex in kiwifruit. Nature plants 5:801–809.

Bachtrog, D. (2013). Y-chromosome evolution: emerging insights into processes of Y-chromosome degeneration. Nature reviews Genetics 14:113–124.

Bawa, K.S. (1980). Evolution of Dioecy in Flowering Plants. Annual Review of Ecology and Systematics 11:15–39.

Bennett, T., van den Toorn, A., Sanchez-Perez, G.F., Campilho, A., Willemsen, V., Snel, B., and Scheres, B. (2010). SOMBRERO, BEARSKIN1, and BEARSKIN2 regulate root cap maturation in *Arabidopsis*. The Plant cell 22:640–654.

Bergero, R., and Charlesworth, D. (2009). The evolution of restricted recombination in sex chromosomes. Trends in ecology & evolution 24:94–102.

Brautigam, K., Soolanayakanahally, R., Champigny, M., Mansfield, S., Douglas, C., Campbell, M.M., and Cronk, Q. (2017). Sexual epigenetics: gender-specific methylation of a gene in the sex determining region of *Populus balsamifera*. Scientific reports 7:45388.

Chae, E., Bomblies, K., Kim, S.T., Karelina, D., Zaidem, M., Ossowski, S., Martin-Pizarro, C., Laitinen, R.A., Rowan, B.A., Tenenboim, H., et al. (2014). Species-wide genetic incompatibility analysis identifies immune genes as hot spots of deleterious epistasis. Cell 159:1341–1351.

Charlesworth, B. (1991). The evolution of sex chromosomes. Science 251:1030–1033.

Charlesworth, B., and Charlesworth, D. (1978). A Model for the Evolution of Dioecy and Gynodioecy. The American Naturalist 112:975–997.

Charlesworth, D. (2012). Plant sex chromosome evolution. Journal of Experimental Botany 64:405–420.

Charlesworth, D. (2016). Plant Sex Chromosomes. Annual Review of Plant Biology 67:397–420.

Chen, Y., Wang, T., Fang, L., Li, X., and Yin, T. (2016). Confirmation of Single-Locus Sex Determination and Female Heterogamety in Willow Based on Linkage Analysis. PloS one 11:e0147671.

Chen, Z., Ai, F., Zhang, J., Ma, X., Yang, W., Wang, W., Su, Y., Wang, M., Yang, Y., Mao, K., et al. (2020) Survival in the Tropics despite isolation, inbreeding and asexual reproduction: insights from the genome of the world’s southernmost poplar (*Populus ilicifolia*). The Plant Journal doi:10.1111/tpj.14744.

Dai, X., Hu, Q., Cai, Q., Feng, K., Ye, N., Tuskan, G.A., Milne, R., Chen, Y., Wan, Z., Wang, Z., et al. (2014). The willow genome and divergent evolution from poplar after the common genome duplication. Cell Research 24:1274–1277.

DePristo, M.A., Banks, E., Poplin, R., Garimella, K.V., Maguire, J.R., Hartl, C., Philippakis, A.A., del Angel, G., Rivas, M.A., Hanna, M., et al. (2011). A framework for variation discovery and genotyping using next-generation DNA sequencing data. Nature genetics 43:491–498.

Feng, G., Sanderson, B.J., Keefover-Ring, K., Liu, J., Ma, T., Yin, T., Smart, L.B., DiFazio, S.P., and Olson, M.S. (2020). Pathways to sex determination in plants: how many roads lead to Rome? Current Opinion in Plant Biology 54:61–68.

Gaudet, M., Jorge, V., Paolucci, I., Beritognolo, I., Mugnozza, G.S., and Sabatti, M. (2008). Genetic linkage maps of *Populus nigra* L. including AFLPs, SSRs, SNPs, and sex trait. Tree Genetics & Genomes 4:25–36.

Geraldes, A., Hefer, C.A., Capron, A., Kolosova, N., Martinez-Nuñez, F., Soolanayakanahally, R.Y., Stanton, B., Guy, R.D., Mansfield, S.D., Douglas, C.J., et al. (2015). Recent Y chromosome divergence despite ancient origin of dioecy in poplars (*Populus*). Molecular Ecology 24:3243–3256.

Gong, L., Wong, C.H., Idol, J., Ngan, C.Y., and Wei, C.L. (2019). Ultra-long Read Sequencing for Whole Genomic DNA Analysis. Journal of visualized experiments: JoVE. 10.3791/58954.

Han, Y., and Jiao, Y. (2015). APETALA1 establishes determinate floral meristem through regulating cytokinins homeostasis in *Arabidopsis*. Plant signaling & behavior 10:e989039.

Harkess, A., Zhou, J., Xu, C., Bowers, J.E., Van der Hulst, R., Ayyampalayam, S., Mercati, F., Riccardi, P., McKain, M.R., Kakrana, A., et al. (2017). The asparagus genome sheds light on the origin and evolution of a young Y chromosome. Nature communications 8:1279.

Henry, I.M., Akagi, T., Tao, R., and Comai, L. (2018). One Hundred Ways to Invent the Sexes: Theoretical and Observed Paths to Dioecy in Plants. Annual Review of Plant Biology 69:553–575.

Hobza, R., Cegan, R., Jesionek, W., Kejnovsky, E., Vyskot, B., and Kubat, Z. (2017). Impact of repetitive elements on the Y chromosome formation in plants. Genes 8:302.

Hobza, R., Kubat, Z., Cegan, R., Jesionek, W., Vyskot, B., and Kejnovsky, E. (2015). Impact of repetitive DNA on sex chromosome evolution in plants. Chromosome Research 23:561–570.

Hou, J., Ye, N., Zhang, D., Chen, Y., Fang, L., Dai, X., and Yin, T. (2015). Different autosomes evolved into sex chromosomes in the sister genera of *Salix* and *Populus*. Scientific reports 5:9076.

Hu, H., Yang, W., Zheng, Z., Niu, Z., Yang, Y., Wan, D., Liu, J., and Ma, T. (2020). Analysis of Alternative Splicing and Alternative Polyadenylation in *Populus alba* var. *pyramidalis* by Single-Molecular Long-Read Sequencing. Frontiers in genetics 11:48.

Jain, M., Koren, S., Miga, K.H., Quick, J., Rand, A.C., Sasani, T.A., Tyson, J.R., Beggs, A.D., Dilthey, A.T., Fiddes, I.T., et al. (2018). Nanopore sequencing and assembly of a human genome with ultra-long reads. Nature Biotechnology 36:338–345.

Kapitonov, V.V., and Jurka, J. (2007). Helitrons on a roll: eukaryotic rolling-circle transposons. Trends in genetics: TIG 23:521–529.

Kersten, B., Pakull, B., Groppe, K., Lueneburg, J., and Fladung, M. (2014). The sex-linked region in *Populus tremuloides* Turesson 141 corresponds to a pericentromeric region of about two million base pairs on *P. trichocarpa* chromosome 19. Plant biology 16:411–418.

Khan, A., Garbelli, A., Grossi, S., Florentin, A., Batelli, G., Acuna, T., Zolla, G., Kaye, Y., Paul, L.K., Zhu, J.K., et al. (2014). The *Arabidopsis* STRESS RESPONSE SUPPRESSOR DEAD-box RNA helicases are nucleolar- and chromocenter-localized proteins that undergo stress-mediated relocalization and are involved in epigenetic gene silencing. The Plant journal 79:28–43.

Kieber, J.J., and Schaller, G.E. (2018). Cytokinin signaling in plant development. Development 145.

Kiełbasa, S.M., Wan, R., Sato, K., Horton, P., and Frith, M.C. (2011). Adaptive seeds tame genomic sequence comparison. Genome research 21:487–493.

Koren, S., Walenz, B.P., Berlin, K., Miller, J.R., Bergman, N.H., and Phillippy, A.M. (2017). Canu: scalable and accurate long-read assembly via adaptive k-mer weighting and repeat separation. Genome research 27:722–736.

Kuroha, T., Tokunaga, H., Kojima, M., Ueda, N., Ishida, T., Nagawa, S., Fukuda, H., Sugimoto, K., and Sakakibara, H. (2009). Functional analyses of LONELY GUY cytokinin-activating enzymes reveal the importance of the direct activation pathway in *Arabidopsis*. The Plant cell 21:3152–3169.

Li, H. (2018). Minimap2: pairwise alignment for nucleotide sequences. Bioinformatics 34:3094–3100.

Li, H., and Durbin, R. (2009). Fast and accurate short read alignment with Burrows-Wheeler transform. Bioinformatics 25:1754–1760.

Li, H., Handsaker, B., Wysoker, A., Fennell, T., Ruan, J., Homer, N., Marth, G., Abecasis, G., and Durbin, R. (2009). The Sequence Alignment/Map format and SAMtools. Bioinformatics 25:2078–2079.

Li, M., Wang, D., Zhang, L., Kang, M., Lu, Z., Zhu, R., Mao, X., Xi, Z., and Ma, T. (2019). Intergeneric Relationships within the Family Salicaceae *s.l.* Based on Plastid Phylogenomics. International journal of molecular sciences 20:3788.

Ma, J., Wan, D., Duan, B., Bai, X., Bai, Q., Chen, N., and Ma, T. (2019). Genome sequence and genetic transformation of a widely distributed and cultivated poplar. Plant biotechnology journal 17:451–460.

Ma, T., Wang, K., Hu, Q., Xi, Z., Wan, D., Wang, Q., Feng, J., Jiang, D., Ahani, H., Abbott, R.J., et al. (2018). Ancient polymorphisms and divergence hitchhiking contribute to genomic islands of divergence within a poplar species complex. Proceedings of the National Academy of Sciences of the United States of America 115:E236–e243.

Melnikova, N.V., Kudryavtseva, A.V., Borkhert, E.V., Pushkova, E.N., Fedorova, M.S., Snezhkina, A.V., Krasnov, G.S., and Dmitriev, A.A. (2019). Sex-specific polymorphism of *MET1* and *ARR17* genes in *Populus* × *sibirica*. Biochimie 162:26–32.

Ming, R., Bendahmane, A., and Renner, S.S. (2011). Sex chromosomes in land plants. Annual Review of Plant Biology 62:485–514.

Mitsuda, N., Seki, M., Shinozaki, K., and Ohme-Takagi, M. (2005). The NAC transcription factors NST1 and NST2 of *Arabidopsis* regulate secondary wall thickenings and are required for anther dehiscence. The Plant cell 17:2993–3006.

Morgante, M., Brunner, S., Pea, G., Fengler, K., Zuccolo, A., and Rafalski, A. (2005). Gene duplication and exon shuffling by helitron-like transposons generate intraspecies diversity in maize. Nature genetics 37:997–1002.

Pakull, B., Groppe, K., Meyer, M., Markussen, T., and Fladung, M. (2009). Genetic linkage mapping in aspen (*Populus tremula* L. and *Populus tremuloides* Michx.). Tree Genetics & Genomes 5:505–515.

Pakull, B., Kersten, B., Lüneburg, J., and Fladung, M. (2015). A simple PCR-based marker to determine sex in aspen. Plant biology 17:256–261.

Palmer, D.H., Rogers, T.F., Dean, R., and Wright, A.E. (2019). How to identify sex chromosomes and their turnover. Molecular Ecology 28:4709–4724.

Paolucci, I., Gaudet, M., Jorge, V., Beritognolo, I., Terzoli, S., Kuzminsky, E., Muleo, R., Scarascia Mugnozza, G., and Sabatti, M. (2010). Genetic linkage maps of *Populus alba* L. and comparative mapping analysis of sex determination across *Populus* species. Tree Genetics & Genomes 6:863–875.

Peto, F. (2011). Cytology of poplar species and natural hybrids. Canadian Journal of Research 16:445–455.

Pucholt, P., Rönnberg-Wästljung, A.C., and Berlin, S. (2015). Single locus sex determination and female heterogamety in the basket willow (*Salix viminalis* L.). Heredity 114:575–583.

Pucholt, P., Wright, A.E., Conze, L.L., Mank, J.E., and Berlin, S. (2017). Recent Sex Chromosome Divergence despite Ancient Dioecy in the Willow *Salix viminalis*. Molecular Biology and Evolution 34:1991–2001.

Purcell, S., Neale, B., Todd-Brown, K., Thomas, L., Ferreira, M.A., Bender, D., Maller, J., Sklar, P., de Bakker, P.I., Daly, M.J., et al. (2007). PLINK: a tool set for whole-genome association and population-based linkage analyses. American journal of human genetics 81:559–575.

Renner, S.S. (2014). The relative and absolute frequencies of angiosperm sexual systems: dioecy, monoecy, gynodioecy, and an updated online database. American journal of botany 101:1588–1596.

Renner, S.S., and Ricklefs, R.E. (1995). Dioecy and its correlates in the flowering plants. American journal of botany:596–606.

Sanderson, B., Feng, G., Hu, N., Grady, J., Carlson, C., Smart, B., Keefover-Ring, K., Yin, T., Ma, T., Liu, J., DiFazio, S., et al. (2020). Sex determination through X-Y heterogamety in *Salix nigra*. bioRxiv.

Stanke, M., Keller, O., Gunduz, I., Hayes, A., Waack, S., and Morgenstern, B. (2006). AUGUSTUS: ab initio prediction of alternative transcripts. Nucleic acids research 34:W435–439.

Tamura, K., Peterson, D., Peterson, N., Stecher, G., Nei, M., and Kumar, S. (2011). MEGA5: molecular evolutionary genetics analysis using maximum likelihood, evolutionary distance, and maximum parsimony methods. Molecular Biology and Evolution 28:2731–2739.

Tarailo-Graovac, M., and Chen, N. (2009). Using RepeatMasker to identify repetitive elements in genomic sequences. Current protocols in bioinformatics Chapter 4:Unit 4.10.

Tennessen, J.A., Wei, N., Straub, S.C.K., Govindarajulu, R., Liston, A., and Ashman, T.L. (2018). Repeated translocation of a gene cassette drives sex-chromosome turnover in strawberries. PLoS biology 16:e2006062.

Thévenin, J., Pollet, B., Letarnec, B., Saulnier, L., Gissot, L., Maia-Grondard, A., Lapierre, C., and Jouanin, L. (2011). The simultaneous repression of CCR and CAD, two enzymes of the lignin biosynthetic pathway, results in sterility and dwarfism in *Arabidopsis thaliana*. Molecular plant 4:70–82.

Tokunaga, H., Kojima, M., Kuroha, T., Ishida, T., Sugimoto, K., Kiba, T., and Sakakibara, H. (2012). *Arabidopsis* lonely guy (LOG) multiple mutants reveal a central role of the LOG-dependent pathway in cytokinin activation. The Plant journal: 69:355–365.

Torres, M.F., Mathew, L.S., Ahmed, I., Al-Azwani, I.K., Krueger, R., Rivera-Nuñez, D., Mohamoud, Y.A., Clark, A.G., Suhre, K., and Malek, J.A. (2018). Genus-wide sequencing supports a two-locus model for sex-determination in *Phoenix*. Nature communications 9:3969.

Tuskan, G.A., and Difazio, S., and Jansson, S., and Bohlmann, J., and Grigoriev, I., and Hellsten, U., and Putnam, N., and Ralph, S., and Rombauts, S., and Salamov, A., et al. (2006). The genome of black cottonwood, *Populus trichocarpa* (Torr. & Gray). Science 313:1596–1604.

Wang, J., Na, J.K., Yu, Q., Gschwend, A.R., Han, J., Zeng, F., Aryal, R., VanBuren, R., Murray, J.E., Zhang, W., et al. (2012). Sequencing papaya X and Y^h^ chromosomes reveals molecular basis of incipient sex chromosome evolution. Proceedings of the National Academy of Sciences of the United States of America 109:13710–13715.

Wang, M., Zhang, L., Zhang, Z., Li, M., Wang, D., Zhang, X., Xi, Z., Keefover-Ring, K., Smart, L.B., DiFazio, S.P., et al. (2020). Phylogenomics of the genus *Populus* reveals extensive interspecific gene flow and balancing selection. The New phytologist 225:1370–1382.

Wybouw, B., and De Rybel, B. (2019). Cytokinin – A Developing Story. Trends in plant science 24:177–185.

Xue, L., Wu, H., Chen, Y., Li, X., Hou, J., Lu, J., Wei, S., Dai, X., Olson, M., Liu, J. (2020). Two antagonistic effect genes mediate separation of sexes in a fully dioecious plant. bioRxiv.

Yang, W., Wang, K., Zhang, J., Ma, J., Liu, J., and Ma, T. (2017). The draft genome sequence of a desert tree *Populus pruinosa*. GigaScience 6:1–7.

Yang, Y., Ma, T., Wang, Z., Lu, Z., Li, Y., Fu, C., Chen, X., Zhao, M., Olson, M.S., and Liu, J. (2018). Genomic effects of population collapse in a critically endangered ironwood tree *Ostrya rehderiana*. Nature communications 9:5449.

Yin, T., Difazio, S.P., Gunter, L.E., Zhang, X., Sewell, M.M., Woolbright, S.A., Allan, G.J., Kelleher, C.T., Douglas, C.J., Wang, M., et al. (2008). Genome structure and emerging evidence of an incipient sex chromosome in *Populus*. Genome research 18:422–430.

Zhang, J., Boualem, A., Bendahmane, A., and Ming, R. (2014). Genomics of sex determination. Current Opinion in Plant Biology 18:110–116.

Zhang, L., Xi, Z., Wang, M., Guo, X., and Ma, T. (2018). Plastome phylogeny and lineage diversification of Salicaceae with focus on poplars and willows. Ecology and evolution 8:7817–7823.

Zhang, Z., Chen, Y., Zhang, J., Ma, X., Li, Y., Li, M., Wang, D., Kang, M., Wu, H., Yang, Y., et al. (2020). Improved genome assembly provides new insights into genome evolution in a desert poplar (*Populus euphratica*). Molecular ecology resources.

Zhou, R., Macaya-Sanz, D., Carlson, C.H., Schmutz, J., Jenkins, J.W., Kudrna, D., Sharma, A., Sandor, L., Shu, S., Barry, K., et al. (2020). A willow sex chromosome reveals convergent evolution of complex palindromic repeats. Genome biology 21:38.

Zhou, R., Macaya-Sanz, D., Rodgers-Melnick, E., Carlson, C.H., Gouker, F.E., Evans, L.M., Schmutz, J., Jenkins, J.W., Yan, J., Tuskan, G.A., et al. (2018). Characterization of a large sex determination region in *Salix purpurea* L. (Salicaceae). Molecular genetics and genomics: MGG 293:1437–1452.

