## Supplemental Figures for "A general model to explain repeated turnovers of sex determination in the Salicaceae"

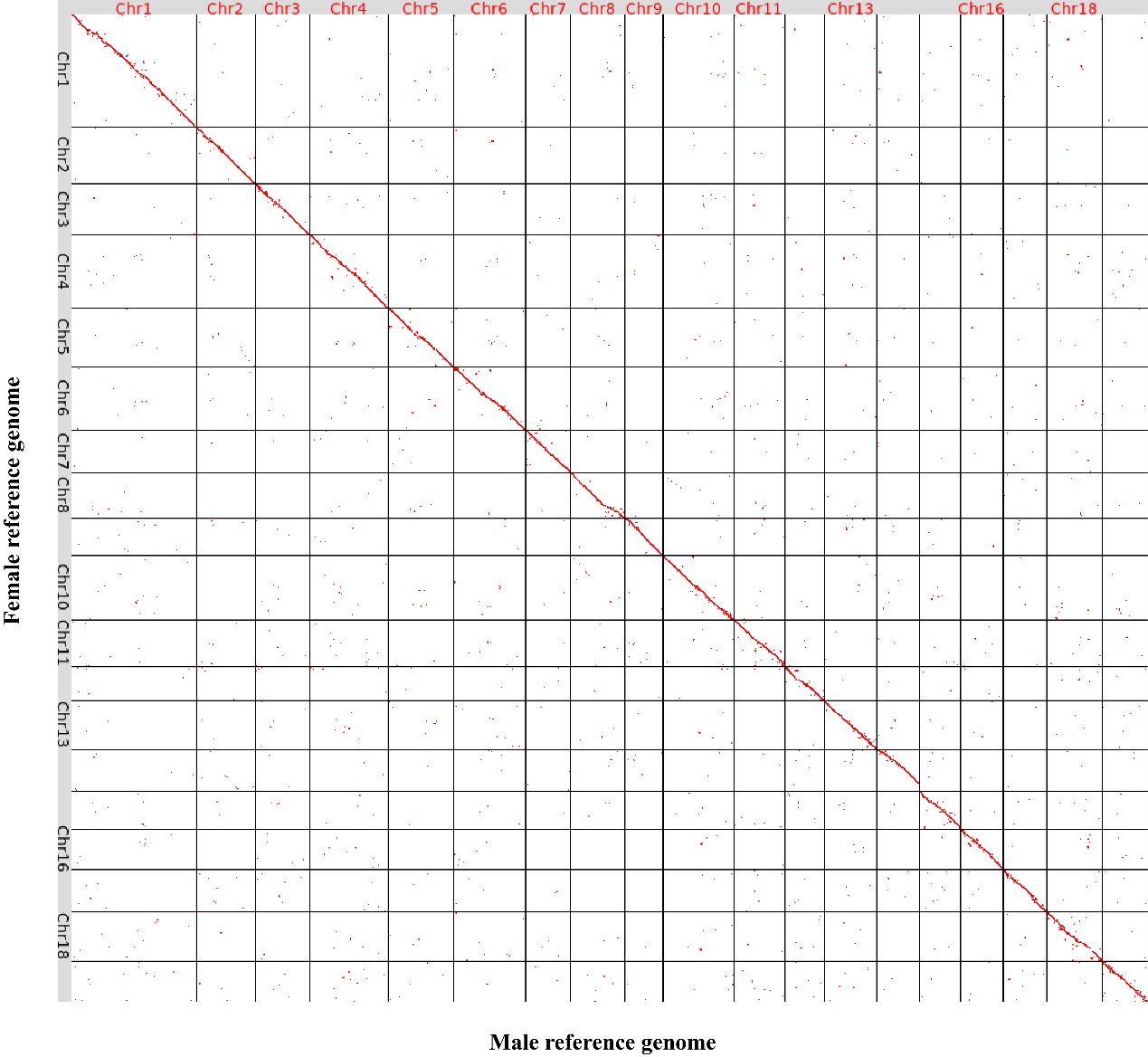

**Fig. S1** **Dot plots of synteny between male and female genome assembly of *P. euphratica*.** The chromosome identities were designated based on alignments with *P. trichocarpa*.

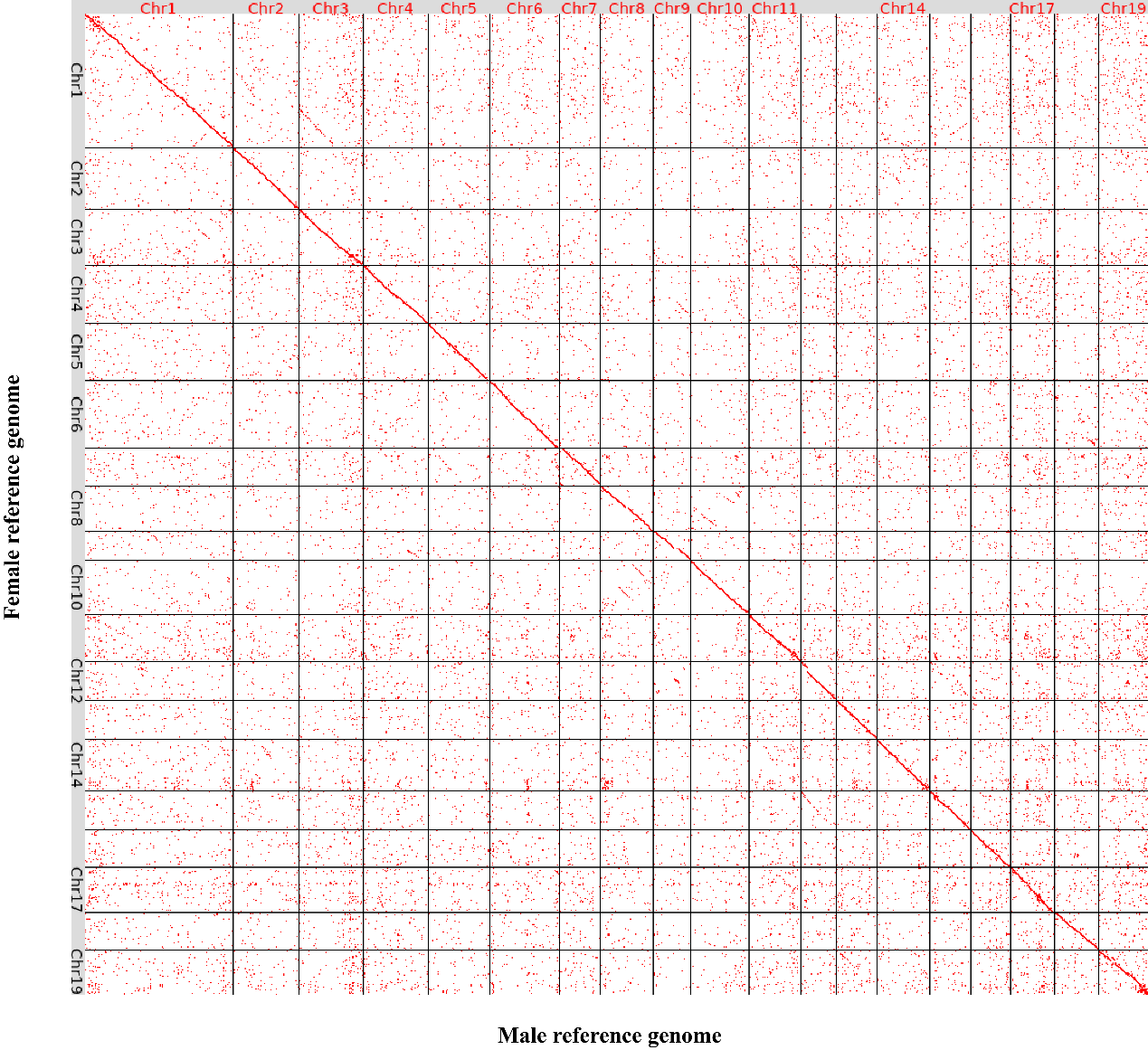

**Fig. S2 Dot plots of synteny between male and female genome assembly of *P. alba*.** The chromosome identities were designated based on alignments with *P. trichocarpa*.

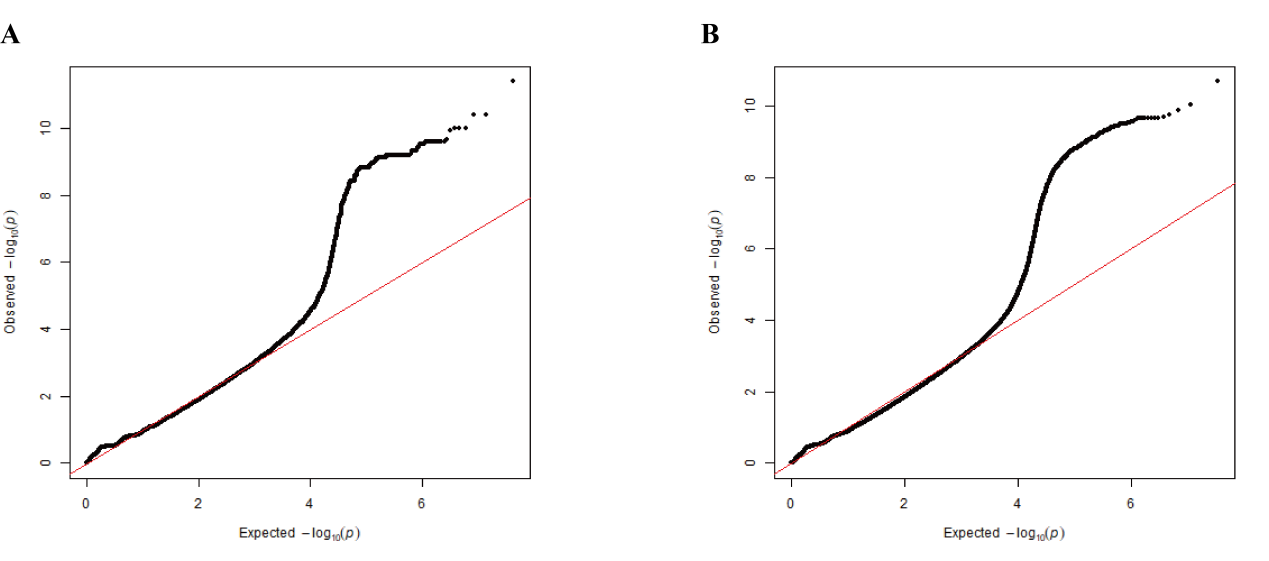

**Fig. S3 Quantile-quantile plot (QQ-plot) derived from genome-wide association study with male (A) and female (B) *P. euphratica* genome as reference, respectively.**

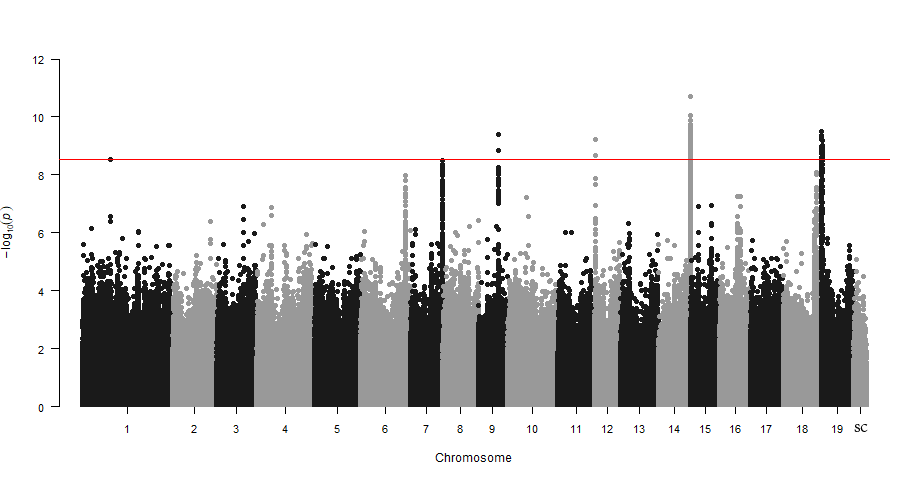

**Fig. S4 Manhattan plot of *P. euphratica* based on the results of genome-wide association study (GWAS) with the female genome as reference.** The y-axis represents the strength of association (−log_10_(*P* value)) for each SNP sorted by chromosomes and scaffolds (SC; x-axis). The red line indicates the significance after Bonferroni multiple corrections (α < 0.05).

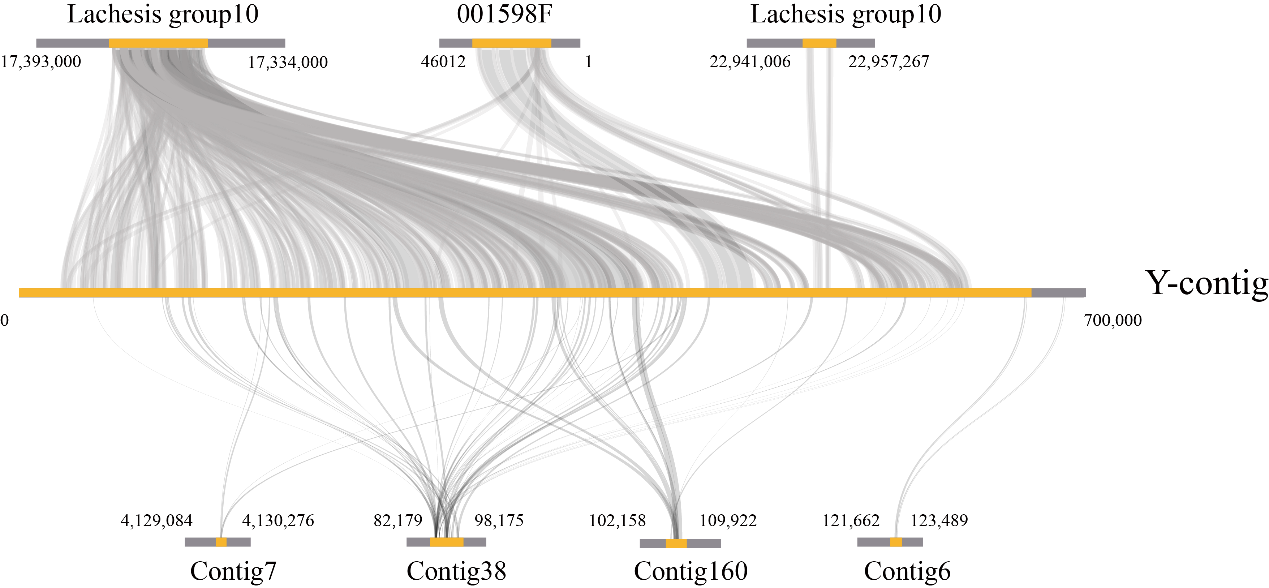

**Fig. S5 High similarity between our assembled Y-contig (highlight in yellow) and the sex-associated regions identified by GWAS using male (top) and female (bottom) *P. euphratica* genome as reference, respectively.** All sex-associated regions contained significant SNPs based on genome-wide association analysis. Detailed information of these sex-associated regions was listed in Fig. 1B and Table S5.

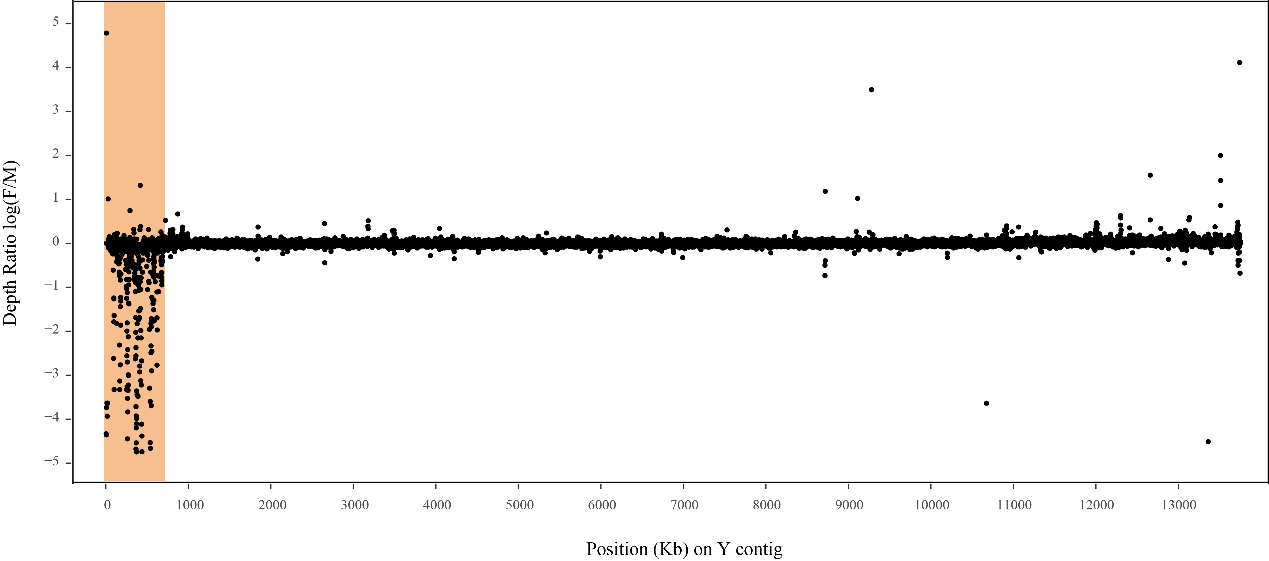

**Fig. S6** **Male-specific depth of contig containing Y-linked region for *P. euphratica*.** Sex depth ratio (log2(F/M)) for 30 female and 30 male individuals are shown by black dots. The Y-linked region is shaded.

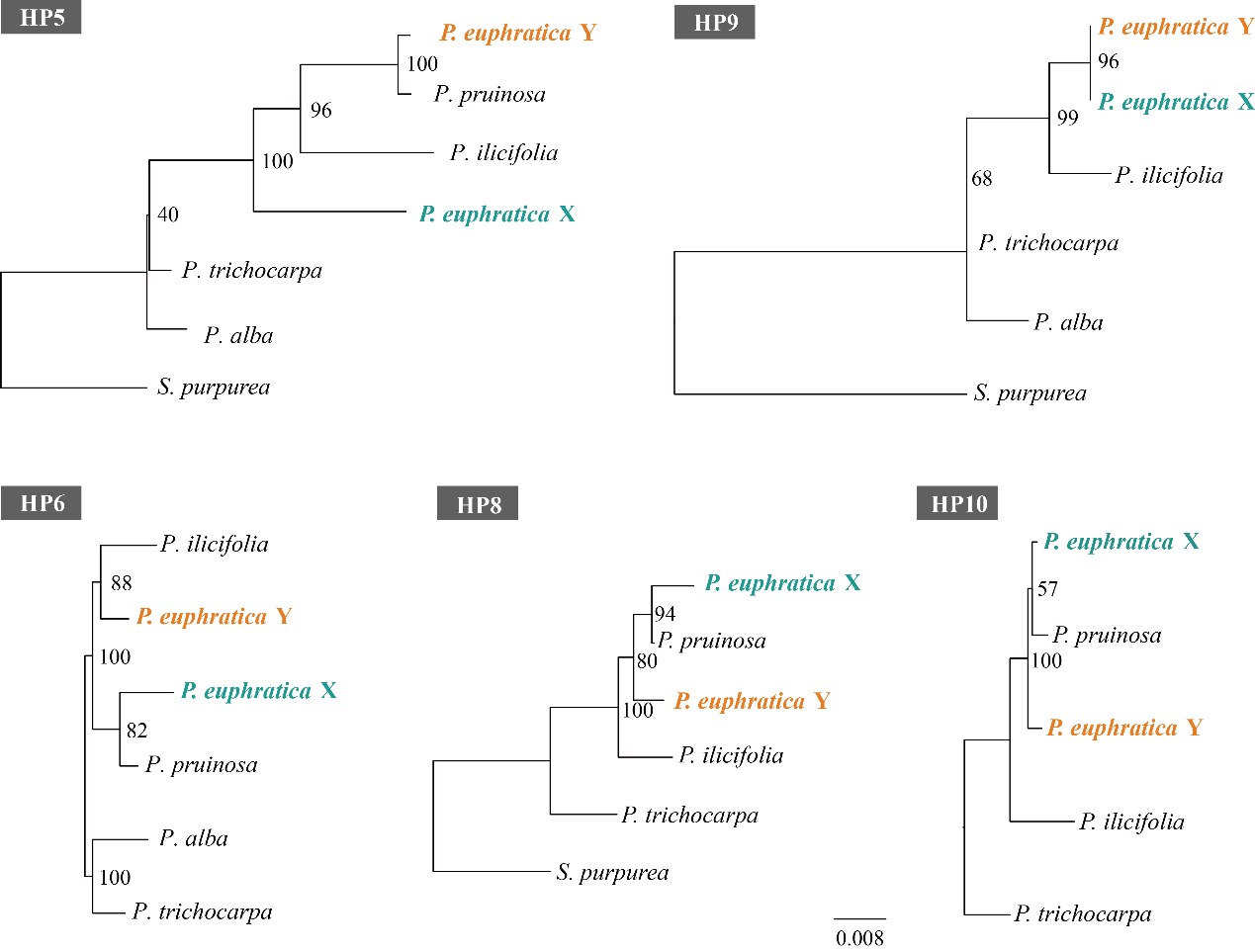

**Fig. S7 Phylogenetic relationships of homologous pairs (HP) shared between Y- and X-SDR of *P. euphratica* and their orthologous genes in other Salicaceae species.** Detailed information about these genes is listed in Table S7 and additional phylogenetic trees are showed in Fig.1D. Note that only orthologous genes located on the corresponding region of chromosome 14 were used for phylogenetic analysis.

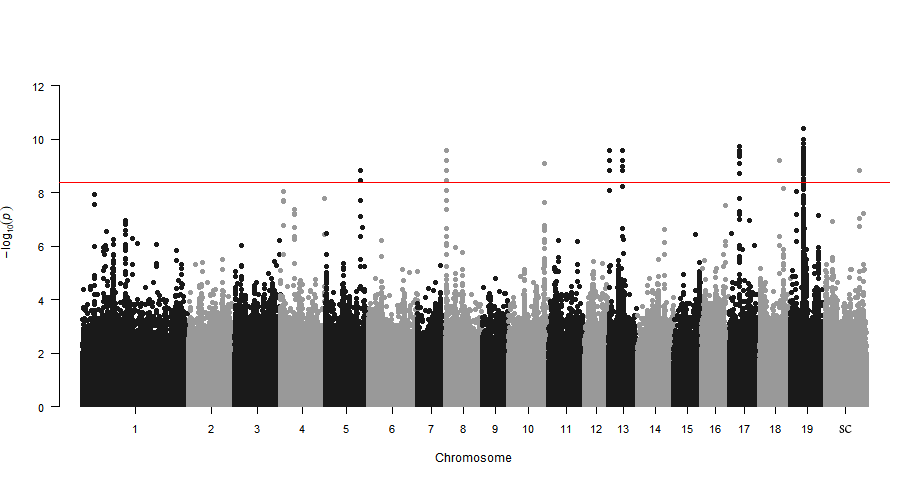

**Fig. S8 Manhattan plot of *P. alba* based on the results of GWAS with the male genome as reference.** The y-axis represents the strength of association (−log_10_(*P* value)) for each SNP sorted by chromosomes and scaffolds (SC; x-axis). The red line indicates the significance after Bonferroni multiple corrections (α < 0.05).

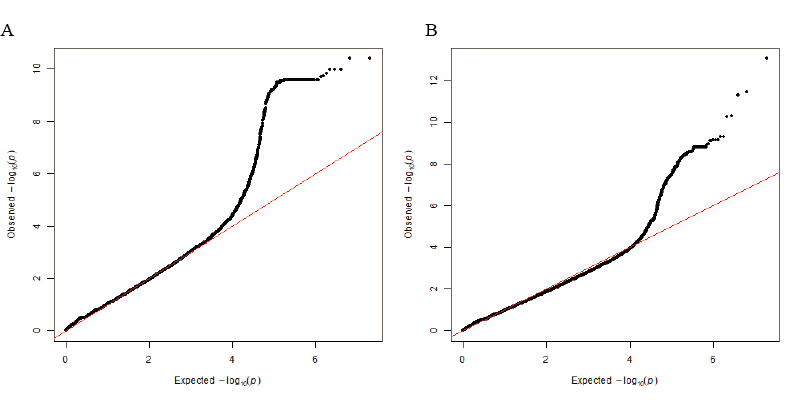

**Fig. S9 Quantile-quantile plot (QQ-plot) derived from genome-wide association study with male (A) and female (B) *P. alba* genome as reference, respectively.**

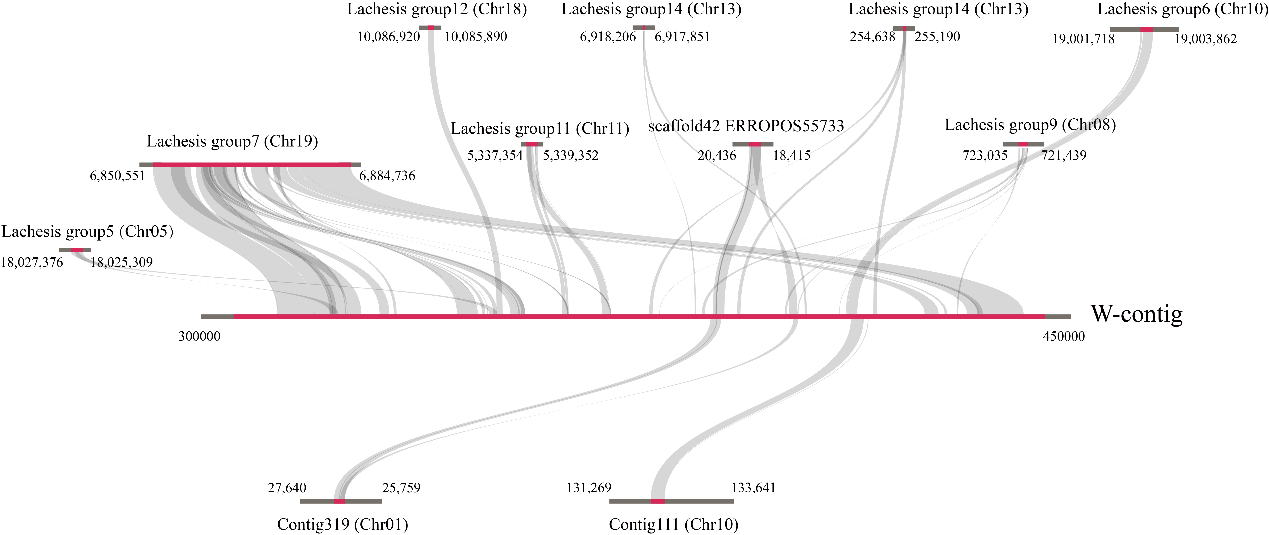

**Fig. S10 High similarity between the assembled W-contig (highlighted in red) and the sex-associated regions identified by GWAS using male (top) and female (bottom) *P. alba* genome as reference.** All of the sex-associated regions contained significant SNPs based on the genome-wide association analysis. Detailed information of these sex-associated regions was listed in Fig. 2B and Table S10.

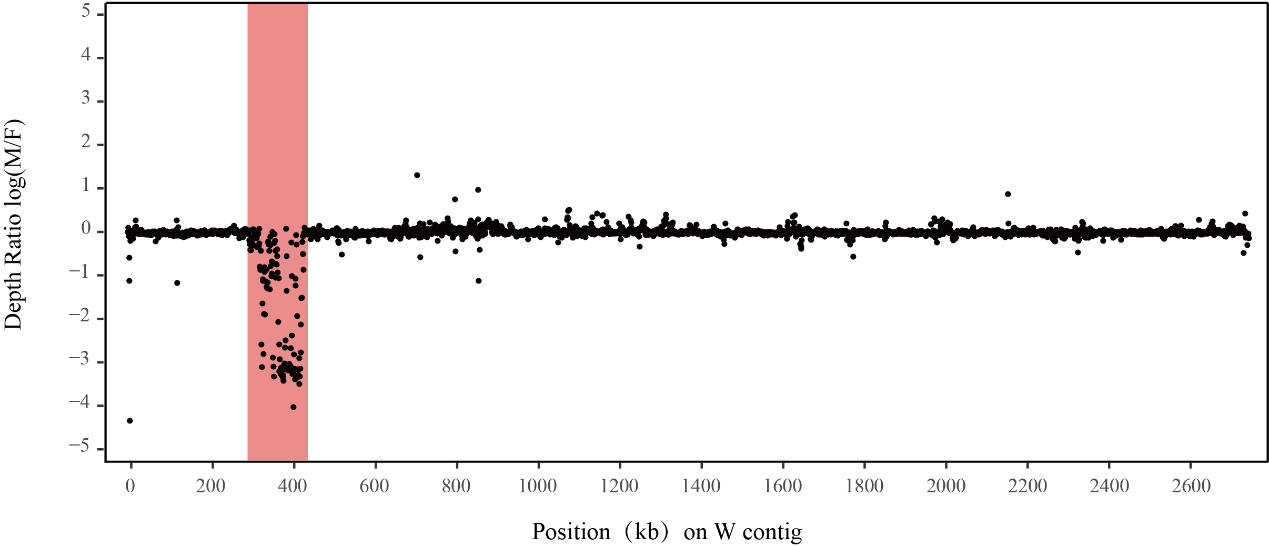

**Fig. S11 Female-specific depth of contig containing W-linked region for *P. alba*.** Sex depth ratio (log2(M/F)) for 30 female and 30 male individuals are shown by black dots. The W-linked region is shaded.
